# Circadian clock disruption promotes the degeneration of dopaminergic neurons

**DOI:** 10.1101/2022.10.15.512376

**Authors:** Michaëla Majcin Dorcikova, Lou C. Duret, Emma Pottié, Emi Nagoshi

## Abstract

Sleep and circadian rhythm disruptions are frequent comorbidities of Parkinson’s disease (PD), a disorder characterized by the progressive loss of dopaminergic (DA) neurons in the substantia nigra. Although sleep/circadian disturbances can be observed years before diagnosing PD, it remains unclear whether circadian clocks have a causal role in the degenerative process. We demonstrated here that circadian clocks regulate the rhythmicity and magnitude of the vulnerability of DA neurons to oxidative stress in *Drosophila*. Circadian pacemaker neurons are presynaptic to a subset of DA neurons and rhythmically modulate their susceptibility to degeneration. The arrhythmic *period* (*per*) gene null mutation exacerbates the age-dependent loss of DA neurons and, in combination with brief oxidative stress, causes premature animal death. These findings suggest that circadian clock disruption promotes dopaminergic neurodegeneration.

## Introduction

Disturbances in the sleep-wake rhythm are well-described symptoms in neurodegenerative disorders such as Parkinson’s disease (PD)^1^. PD is a complex disorder characterized by the progressive loss of dopaminergic (DA) neurons in the substantia nigra pars compacta, leading to motor symptoms^2^. Sleep and circadian disruption can be observed years before the onset of motor symptoms^3^, raising the possibility that circadian rhythm disruption precedes and contributes to PD development. Studies using animal models of PD showed that circadian disruption could exacerbate neurodegeneration and motor symptoms^4^. However, it remains elusive whether circadian disturbances causally drive neurodegeneration or they are caused by neurodegeneration, partly because PD pathology includes dysfunction of the brain regions regulating sleep, resulting in disrupted sleep-wake cycles^5^. Dissecting mechanistic links between circadian clock disruption and neurodegeneration—two intertwined and potentially synergistic processes—is critical for developing reliable diagnostic tools and treatment strategies for PD.

*Drosophila melanogaster* is a genetically tractable model system for studying neurodegenerative diseases^6^. Taking advantage of the conservation of PD-related molecular genetic pathways, numerous genetic and toxin-induced models of PD that display varying degrees of neurodegeneration and motor and non-motor symptoms have been generated^7,8^. The adult fly midbrain contains approximately eight clusters of DA neurons projecting to various brain regions^9^. Functional impairments or loss of DA neurons in the protocerebral anterior medial (PAM) and protocerebral anterior lateral 1 (PPL1) clusters have been consistently observed in PD models, in addition to the losses in other clusters^10–13^. The PAM cluster comprises ∼130 neurons per hemisphere, accounting for approximately 80% of fly brain DA neurons. PAM neurons are highly heterogeneous in their projection patterns and functions and play critical roles in behavioral processes such as olfactory associative learning, foraging, sleep, and locomotion^14–18^. We and others showed that loss of a subclass of PAM neurons leads to defective startle-induced climbing behavior in several PD models^13,19,20^, suggesting a partial analogy between the PAM cluster and the substantia nigra pars compacta.

Circadian clocks, built upon the conserved design principle of negative feedback loops, drive rhythms in numerous behavioral and physiological processes in species across phylogenic trees. The core feedback loop of the *Drosophila* circadian clock consists of CLOCK/CYCLE (CLK/CYC) heterodimers that activate transcription of the *period* (*per*) and *timeless* (*tim*) genes and PER and TIM proteins that feedback-inhibit CLK/CYC activity. CLK/CYC also activates the transcription of genes encoding VRILLE (VRI) and PDP-1, negatively and positively regulating *Clk*, forming a stabilizing loop interlocked with the core loop ^21^. Circadian clocks are present in approximately 150 pacemaker neurons and 1800 glia in the fly brain^22^. Pacemaker neurons are classified into functionally and anatomically diverse subgroups: small and large lateral ventral neurons (s- and l-LNvs), lateral dorsal neurons (LNds), lateral posterior neurons, and three groups of dorsal neurons (DN1s, DN2s, and DN3s). The s- and l-LNvs express the neuropeptide pigment-dispersing factor (PDF), essential for synchronizing rhythmicity across the pacemaker circuit. The PDF-positive s-LNvs (the M-cells) drive morning peak of activity under light-dark cycles (LD) and control free-running 24-h period locomotor rhythms in constant darkness (DD). The robustness and flexibility of the locomotor behavior adapting to environmental conditions require the communication between M-cells and the second set of pacemaker subtypes, the E-cells, comprising the PDF-negative fifth s-LNv, half of the LNds, and some of the DN1s (Reviewed in ^23^). The pacemaker circuit controls circadian locomotor output and regulates brain-wide rhythmic physiology by conveying time-of-day information to areas in the brain such as the mushroom body (MB)^24^.

Misalignment between environmental cycles and endogenous circadian rhythms increases the risk for physical and psychiatric disorders^25^. Genetic and environmental disruption of circadian rhythms in flies leads to adverse health consequences, including increased mortality in response to oxidative stress^26^ and lifespan reduction^27^, suggesting that the mechanisms underpinning the link between circadian disruption and diseases can be studied in flies. *Clk* loss of function in the s-LNv pacemaker neurons causes loss of DA neurons in the PPL1 cluster, resulting in age-dependent locomotor decline; intriguingly, this process is PDF-dependent and circadian-clock-independent^28^. Therefore, questions remain regarding whether and how the circadian system regulates the (patho-)physiology of DA neurons.

Here, we used *Drosophila* as a model to ask whether circadian clocks causally impact degenerative processes in PD. We found that DA neurons in the PAM cluster exhibit diurnal and circadian rhythms in vulnerability to oxidative stress. Loss of circadian clock genes results in age-dependent PAM neuron loss and exacerbates oxidative stress-induced PAM neuron loss. We identified PAM-*α*1 neurons as the selectively and rhythmically susceptible DA neuron subtype and found that circadian pacemaker circuit is presynaptic to this subgroup. Oxidative stress-induced loss of PAM-*α*1 neurons leads to changes in sleep, reminiscent of PD non-motor symptoms. These findings establish a direct role of circadian clocks in controlling the timing and magnitude of the vulnerability of DA neurons.

## Results

### Circadian vulnerability of DA neurons to oxidative insults

We previously demonstrated that flies exposed to hydrogen peroxide (H_2_O_2_) or paraquat for 24 h display selective loss of DA neurons in the PAM cluster (Fig. 1A)^12,13,20^. To evaluate dopaminergic neurodegeneration from a circadian perspective, we developed a short-term H_2_O_2_ treatment protocol that can be performed at six timepoints across the day. 7-day-old *w*^*1118*^ flies were treated with H_2_O_2_ at various concentrations (5% to 20%) and durations (3 h to 6 h), and survival of PAM neurons was assessed 7 days post-treatment by immunostaining for tyrosine hydroxylase (TH) (Fig. S1A). From the conditions that induced PAM neuron loss, we chose 5% and 10% H_2_O_2_ treatment for 4 h in subsequent experiments because this duration is suitable for performing assays in a circadian fashion and the effect does not saturate. Loss of TH-positive neurons was observed only in the PAM cluster among all DA neuron clusters following the short H_2_O_2_ treatment (Fig. 1B). H_2_O_2_ treatment in flies expressing red nuclear fluorescent protein RedStinger with the PAM neuron-specific *R58E02* GAL4 driver resulted in the reduction of RedStinger-positive cells, suggesting that oxidative insults cause loss of neurons and not merely the reduction in TH expression (Fig. S1B). Therefore, we focused on this group of DA neurons in subsequent studies.

**Fig. 1.**
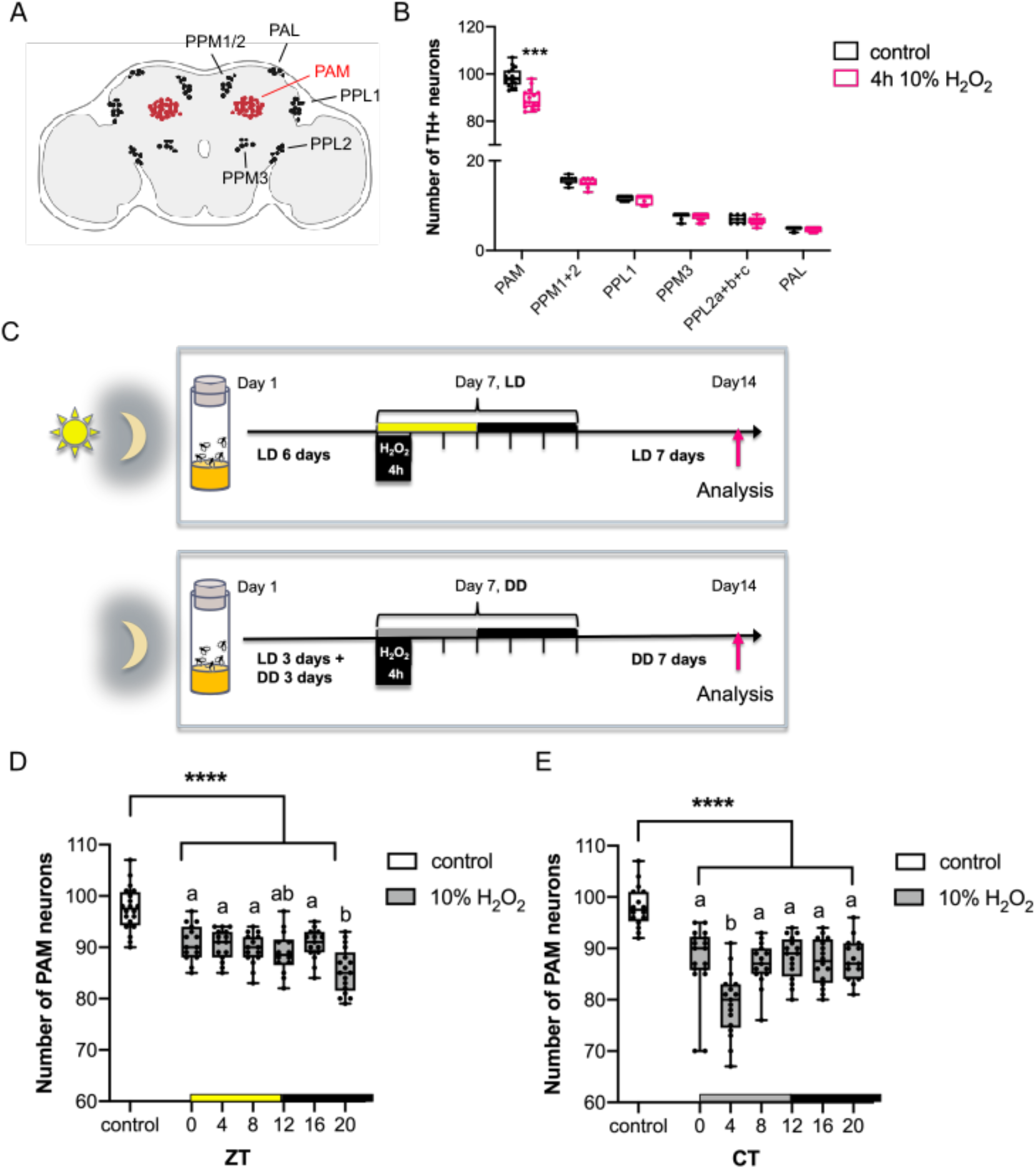
Rhythmic vulnerability of PAM neurons to oxidative stress. (**A**) A schematic of the cell bodies of DA neuron clusters in the adult fly brain. PAM, protocerebral anterior medial. PAL, protocerebral anterior lateral. PPL1 and 2, protocerebral posterior lateral 1 and 2. PPM1, 2, and 3, protocerebral posterior medial 1, 2, and 3. (**B**) DA neuron counts in *w*^*1118*^ flies per hemisphere in each cluster 7 days after a 4-h 10% H_2_O_2_ treatment performed at ZT20. The control group was treated with water only. DA neurons were detected by anti-TH immunostaining. Neurodegeneration was observed only in the PAM cluster following the H_2_O_2_ treatment. \*\*\**p*<0.0001 (t-test, comparing the control and H_2_O_2_-treated groups.). n = 11–18 hemispheres. In box plots in this and all following figures, box boundaries are the 25^th^ and 75^th^ percentiles, the horizontal line across the box is the median, and the whiskers indicate the minimum and maximum values. The dots represent all data points. (**C**) A schematic of the circadian H_2_O_2_ treatment protocol. (**D** and **E**) Quantification of the number of PAM neurons after the 4-h 10% H_2_O_2_ treatment performed at different times in LD (**D**) or DD (**E**). The x-axis indicates the times when H_2_O_2_ was applied. n = 14–20 hemispheres. The control group was treated with water only at ZT20 in LD (**D**) and CT20 in DD (**E**). At all timepoints, PAM neuron counts in the H_2_O_2_ treatment group are significantly smaller than those in the control group. \*\*\*\**p*<0.0001 (ANOVA with Tukey’s post hoc test). Within the H_2_O_2_-treated group, flies treated at ZT20 in LD (**D**) and CT4 in DD (**E**) showed significantly greater cell loss than the treatment at any other time. Different lowercase letters represent statistical significance by ANOVA with Tukey’s HSD post hoc test.

To test whether the vulnerability of PAM neurons to oxidative stress is rhythmic, we next performed the short H_2_O_2_ treatment in a circadian fashion. 7-day-old *w*^*1118*^ flies entrained to LD (12h:12h light-dark cycles) were exposed to 10% H_2_O_2_ for 4 h, following a 5-h food and water deprivation. H_2_O_2_ exposure was initiated at a different Zeitgeber Time (ZT) across the day, and DA neurons were analyzed 7 days post-treatment. (Fig. 1C). Whereas the H_2_O_2_ treatment at any time caused significant PAM neuron loss compared to the water-only control, the treatment at ZT20 (8 h after lights-off) resulted in a more significant loss of PAM neurons than the other timepoints (Fig. 1D). The highest sensitivity at ZT20 was replicated in the assay with the 4-h 5% H_2_O_2_ treatment, validating its specificity (Fig. S1C). We performed the same experiment in constant darkness (DD) to determine the role of endogenous clocks and LD cycles. Newly hatched flies were entrained for three days in LD and then placed in DD for the remainder of the experiment and treated with H_2_O_2_ on the fourth day in DD (DD4) (Fig. 1C). The treatment at CT4 (4 h after subjective lights-on) with 10% (Fig. 1D) or 5% H_2_O_2_ (Fig. S1D) caused a more significant PAM neuron loss than the other timepoints. These results suggest intrinsic rhythmicity in the vulnerability of PAM neurons to oxidative stress modulated by light.

To determine whether circadian clocks control the intrinsic rhythms of PAM neuron vulnerability, we next used *per* null (*per*^*01*^, also known as *per*^*0*^) mutant flies devoid of molecular clocks. We first quantified PAM neuron counts in *per*^*0*^ with anti-TH staining and by expressing RedStinger without oxidative stress from days 1 to 14 of age. We found that *per*^*0*^ flies were born with fewer PAM neurons and displayed a marked, age-dependent loss of PAM neurons (Fig. 2A-C). The *tim* null (*tim*^*0*^) mutants similarly displayed a reduced number of PAM neurons on day 1 and a progressive loss thereafter (Fig. S2A). Both *per*^*0*^ and *tim*^*0*^ arrhythmic mutants displayed locomotor deficits, as reported previously^28,29^ (Fig. S2B). These findings suggest that disruption of circadian clocks interferes with the development of DA neurons in the PAM cluster and accelerates their age-dependent degeneration.

**Fig. 2.**
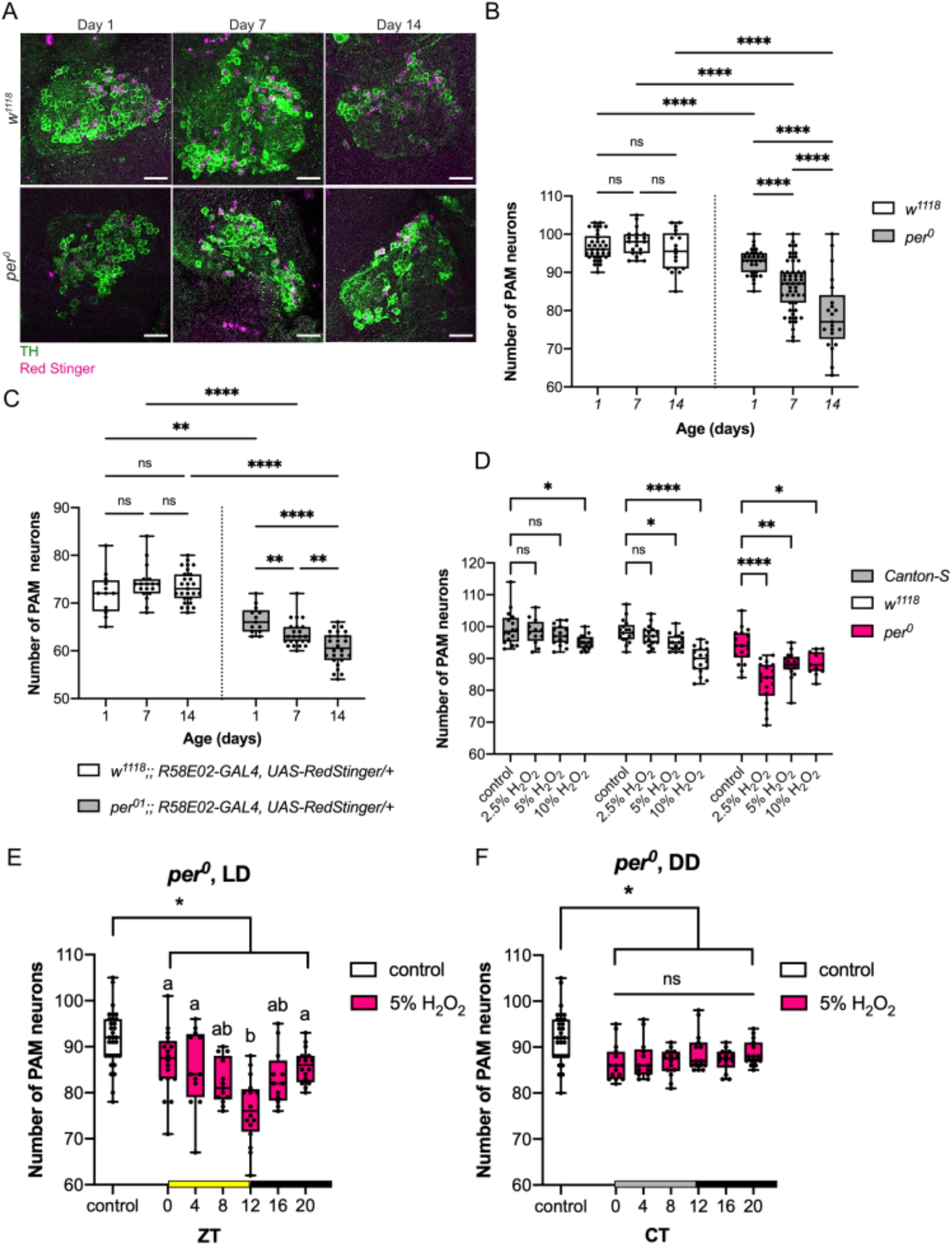
The clock gene *per* controls the magnitude and rhythms of the vulnerability of PAM neurons to oxidative stress. (**A**) Representative images of PAM neurons in *w*^*1118*^ and *per*^*0*^ flies at indicated ages. PAM neurons were visualized by anti-TH antibodies (green) and RedStinger driven by the *R58E02* driver (magenta). Scale bar, 20 *μ*m. (**B** and **C**) The number of PAM neurons detected by anti-TH immunostaining (**B**) and by the expression of *R58E02*-driven RedStinger (**C**). n = 13–26 hemispheres. *per*^*0*^ flies display developmental and age-dependent loss of PAM neurons. \*\**p*<0.01 and \*\*\*\**p*<0.0001 (t-test or Mann-Whitney U test). (**D**) PAM neuron counts in *Canton-S, w*^*1118*^, and *per*^*0*^ were assessed by anti-TH staining 7 days after the treatment with different concentrations of H_2_O_2._ Treatment was performed from ZT1 for 4 h. n = 12-18 hemispheres. *per*^*0*^ flies display increased PAM neuron susceptibility compared control genotypes. \**p*<0.05, \*\*\**p*<0.001, and \*\*\*\**p*<0.0001 (ANOVA with Dunnett’s multiple comparisons test). (**E** and **F**) PAM neuron counts in *per*^*0*^ flies after a 4-h 5% H_2_O_2_ treatment were performed at different timepoints in LD (**E**) or DD (**F**). The x-axis indicates the timepoints when H_2_O_2_ was applied. n = 12–32 hemispheres. The control group was treated with water only at ZT0 in LD (**E**) and CT0 in DD (**F**). At all timepoints, PAM neuron counts in the H_2_O_2_ treatment group were significantly smaller than those in the control group. \**p*<0.05 (ANOVA with Tukey’s HSD post hoc test). Within the H_2_O_2_-treated group, flies treated at ZT12 in LD (**E**) displayed a significantly greater cell loss than at any other timepoint. No difference was observed between timepoints in DD (**F**). Different lowercase letters represent statistical significance by ANOVA with Tukey’s post hoc test.

Because a previous study reported that H_2_O_2_ exposure elicits a higher rate of mortality in *per*^*0*^ than in wild-type flies^29^, we also examined the effect of various concentrations of H_2_O_2_ on PAM neuron survival in *per*^*0*^ flies at a single timepoint; 2.5% H_2_O_2_, which did not affect PAM neurons in *w*^*1118*^ and *Canton-S* (*CS*), robustly induced PAM neurodegeneration in *per*^*0*^ mutants (Fig. 2D). This finding suggests that loss of *per* enhances the vulnerability of PAM neurons to oxidative stress-induced degeneration. We next performed the circadian oxidative treatment with 5% H_2_O_2_ in LD and DD on *per*^*0*^ flies. The *per*^*0*^ flies treated with H_2_O_2_ at any timepoint showed a significant loss of PAM neurons than the control group treated with water only. In LD, the treatment at ZT12 resulted in a more significant loss of PAM neurons than the other timepoints (Fig. 2E). In contrast, the difference among timepoints was abolished in DD (Fig. 2F). These findings suggest that the circadian clock gene *per* controls the magnitude and temporal variations of PAM neuron sensitivity to oxidative stress. In the absence of functional clocks, light can impose diurnal rhythms of PAM neuron vulnerability.

### The circadian neural circuit controls the rhythmic vulnerability of PAM neurons

Because PAM neurons are not clock-containing cells, the rhythmic vulnerability of PAM neurons is non-cell-autonomously controlled by clocks located elsewhere. Several mechanisms might drive rhythms in non-clock-containing neurons, including the rhythmic H_2_O_2_ uptake by feeding/drinking rhythms, systemic rhythmic changes in the brain environment, and the rhythmic modulation of the physiology of PAM neurons by circadian pacemaker neurons.

To test whether the differential loss of PAM neurons is caused by ingesting more H_2_O_2_ solution at one time than the others, we performed a feeding assay^30^. Flies were fed with H_2_O_2_ or water mixed with blue food dye, which sticks to the gut and is not eliminated or digested. Then, spectrophotometry was performed on homogenized fly bodies to measure the amount of dye ingested. In LD and DD, flies displayed rhythmic feeding patterns when fed with the control solution (water and dye), congruent with a previous study demonstrating the circadian regulation of feeding^31^. In contrast, the feeding pattern of the 10% H_2_O_2_-containing solution was not rhythmic. Its uptake was significantly lower than the control solution across all timepoints (Fig. 3A). This finding is consistent with the finding that 2.5% H_2_O_2_ caused a more significant neuronal loss than the higher concentration in *per*^*0*^ flies (Fig. 2D), probably because flies ingest more H_2_O_2_ solutions at lower concentrations. Therefore, daily variations in the feeding pattern do not account for the circadian vulnerability of PAM neurons. We also found that flies drink three to five times more in DD than in LD, which explains the more pronounced loss of PAM neurons following the H_2_O_2_ treatment in DD than in LD (Figs. 1D and E, Figs. S1C and D).

**Fig. 3.**
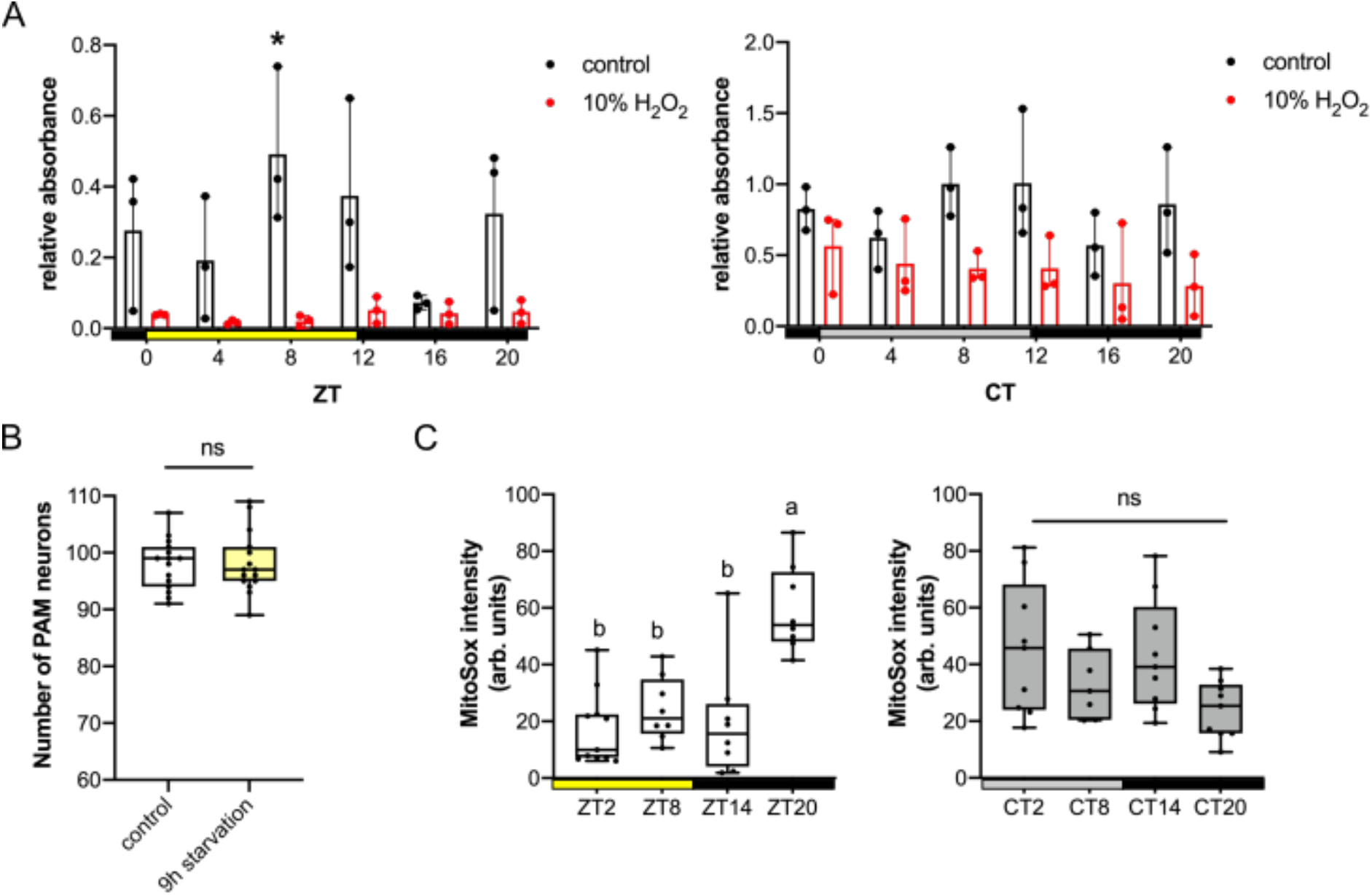
Feeding or dehydration is not the cause of circadian vulnerability of PAM neurons. (**A**) Feeding patterns of *w*^*1118*^ flies in LD (left) and DD (right). Flies were fed for 4 h with 10% H_2_O_2_ solution containing blue dye. The control solution contained only water and blue food dye. n = 15–25 flies. The x-axis indicates the timepoints when the flies started to be fed. The y-axis represents relative absorbance per ten flies. The dots represent the values of three independent experiments. Feeding levels in the control group at ZT8 were significantly higher than at other timepoints in LD. \**p*<0.05 (ANOVA with Tukey’s post hoc test). No significant difference was observed between timepoints in all other groups. (**B**) PAM neuron counts after 9 h of food and water deprivation, started at ZT15. The control group was deprived of food and water for 5 h, followed by 4 h of water access. n = 15–16 hemispheres. No significant difference between groups (t-test). (**C**) ROS levels within the PAM neurons were measured using MitoSOX red in flies expressing *EGFP* with the *R58E02* driver in LD (left) and DD (right). n = 7–11 flies. ROS levels are significantly elevated at ZT20 in LD but do not differ among timepoints in DD (ANOVA with Tukey’s post hoc test).

The circadian H_2_O_2_ feeding assay requires flies to be starved before and during H_2_O_2_ exposure. We next tested whether the starvation and dehydration associated with the assay led to the rhythmic PAM neurodegeneration. To this end, flies were removed from food and water for 9 h, 5 h before and 4 h after ZT20. 7 days later, PAM neurons were examined by anti-TH immune staining. The control group was given access to water for 4 h after a 5-h starvation/water deprivation period. The results showed no changes in PAM neuron count between the groups (Fig. 3B). This finding suggests that the loss of PAM neurons and their rhythmicity are not caused by starvation or dehydration before or during the H_2_O_2_ treatment.

Endogenous oxidative stress levels vary daily in flies and mammals^26,32^. We wondered whether the rhythms in endogenous reactive oxygen species (ROS) levels in the fly brain and H_2_O_2_ administered at specific times would accumulate to trigger a rhythmic neuronal loss in the PAM cluster. To test this possibility, we used MitoSOX (a ROS-sensitive fluorescent dye targeted to mitochondria) to monitor mitochondrial ROS levels across the day in PAM neurons labeled with *R58E02>UAS-EGFP* in *w*^*1118*^ flies. In 7-day-old flies, MitoSOX fluorescence intensity in PAM neurons in LD showed daily variations and peaked at ZT20 (Fig. 3C), the time when PAM neurons are the most vulnerable to H_2_O_2_ (Fig. 1D). This finding suggests that oscillating levels of endogenous ROS combined with the H_2_O_2_ treatment might account for the differential loss of PAM neurons in LD. In contrast, no significant differences in MitoSOX levels were observed between timepoints in DD (Fig. 3C). Therefore, another mechanism under the control of circadian clocks gates the vulnerability of PAM neurons in DD.

To test whether PER-expressing clock neurons directly or indirectly modulate PAM neuron vulnerability, we re-expressed the PER protein in clock neurons in *per*^*0*^ mutants and examined PAM neurons. Genetic rescue of *per* with the pan-clock neuron driver *Clk1982-GAL4*^33^ significantly increased the number of PAM neurons at day 14 compared to *per*^*0*^ without H_2_O_2_ treatment. Thus, the pan clock-neuron *per* rescue prevented the accelerated age-dependent loss of PAM neurons caused by loss of *per* (Fig. 4A). *Clk1982>per* also reversed the exacerbated loss of PAM neurons following H_2_O_2_ treatment in *per*^*0*^ to the level of *w*^*1118*^ flies (Fig. 4B). These results suggest that loss of PER in clock neurons is responsible for the enhanced PAM neuron loss in *per*^*0*^ mutants, at least in part.

**Fig. 4.**
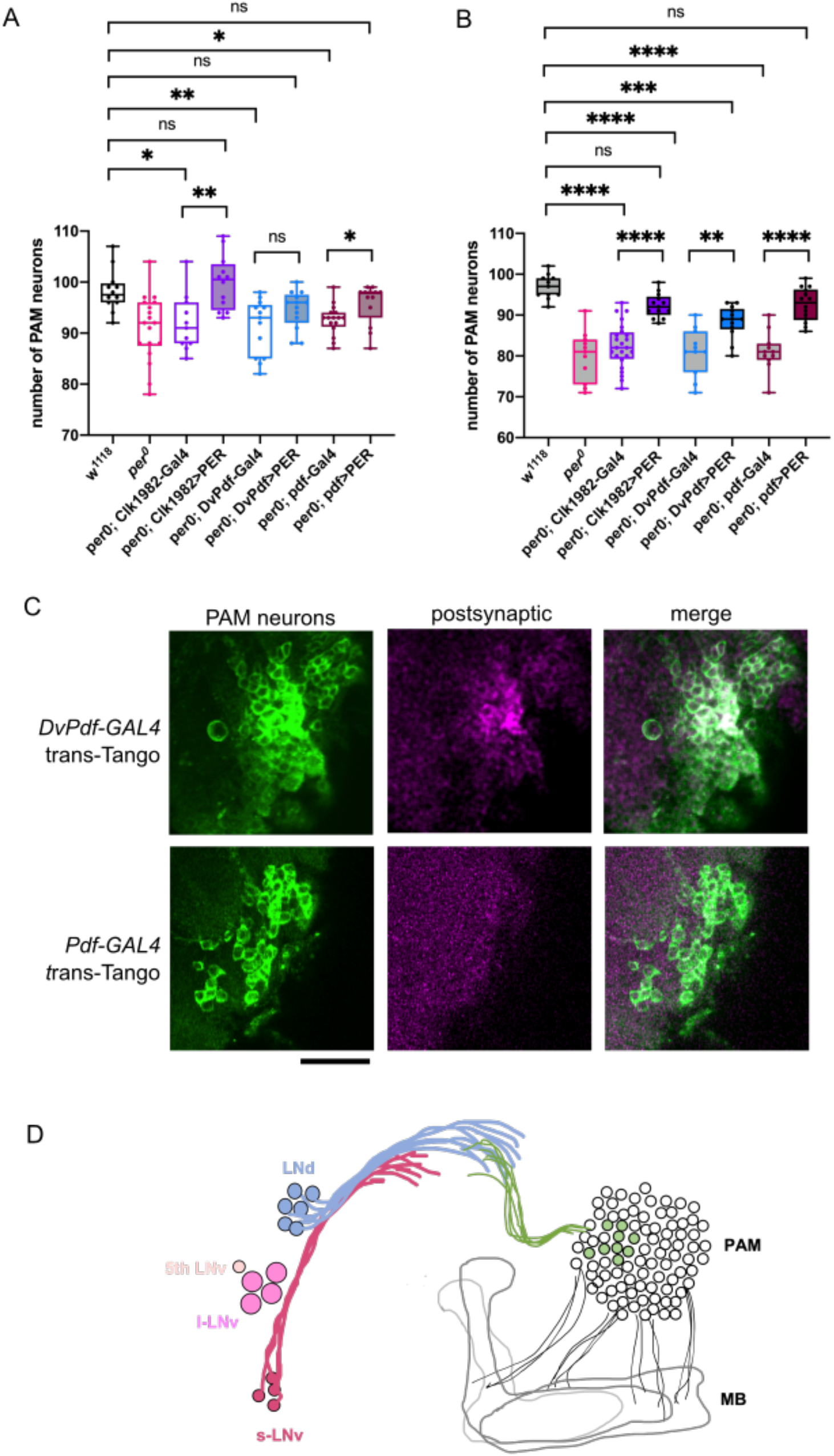
LNd clock neurons are presynaptic to PAM neurons. (**A** and **B**) Genetic rescue of *per* in clock neurons in *per*^*0*^ mutants. PAM neurons were counted using anti-TH staining 7 days after a control treatment with water (**A**), and after a 4-h treatment with 2.5% H_2_O_2_ performed at ZT12 **(B**). Genotypes are indicated on the x-axis. a>b represents that UAS-transgene b is driven by the GAL4 driver a. n = 11–24 hemispheres. Rescue genotypes significantly improve the preservation of PAM neurons after the H_2_O_2_ treatment. \**p*<0.05, \*\**p*<0.01, \*\*\**p*<0.001, and \*\*\*\**p*<0.0001 (ANOVA with Dunnett’s multiple comparisons test). (**C)** *trans*-Tango experiments using *DvPdf-GAL4* and *Pdf-GAL4* combined with TH immunostaining (green). Several postsynaptic targets of *DvPdf* neurons (magenta) were identified as PAM neurons, whereas no postsynaptic signal of *Pdf* neurons was found within the PAM cluster. Presynaptic signals by *DvPdf-GAL4* and *Pdf-GAL* are not visible in these pictures. Scale bar, 20 *μ*m. (**D**) A schematic representation of the *trans*-Tango labeling results. *DvPdf-GAL4*-positive but *Pdf-GAL4*-negative neurons, i.e., LNds, are presynaptic to a subset of PAM neurons.

To test if any subclass of clock neurons is essential for the role of PER in regulating PAM neuron survival, we next performed *per*^*0*^ genetic rescue using *Pdf-GAL4*, expressed in the s- and l-LNvs except the fifth s-LNv ^34^, and *DvPdf-GAL4*, which targets all LNvs, three CRY-negative LNds, and one CRY/ITP double-positive LNd^35,36^. Both rescue genotypes restored the number of PAM neurons to the level of *w*^*1118*^ under basal conditions (Fig. 4A). The number of surviving PAM neurons after H_2_O_2_ treatment was also significantly greater in both rescue genotypes than in the control expressing only the driver (Fig. 4B). PER rescue with *Pdf-GAL4* or *Clk1982-GAL4* showed similar levels of protection against H_2_O_2_, whereas, with *DvPdf-GAL4*, the protection was partial. This might be due to differential expression levels within the LNvs between *Pdf-GAL4* and *DvPdf-GAL4*. Taken together, these findings suggest that PER expression within the PDF-positive LNvs plays a significant role in the regulation of survival of PAM neurons following oxidative insults.

### A subset of clock neurons are presynaptic to PAM neurons

The finding that PER expressed in circadian pacemaker neurons controls rhythmic PAM neuron vulnerability suggests that PAM neurons are directly or indirectly downstream of circadian neural circuits. To test this possibility, we examined postsynaptic targets of *DvPdf-GAL4*-expressing neurons using *trans*-Tango^37^, an anterograde transsynaptic tracing tool. Several neurons in the most anterior region of the PAM cluster were marked with the postsynaptic signal, indicating the direct projection from the *DvPdf-GAL4*-labeled neurons to 6 to 7 PAM neurons (Fig. 4C). In contrast, the *trans*-Tango assay with *Pdf-GAL4* did not result in any positive signal within the PAM cluster (Fig. 4C), suggesting that the s- and l-LNv neurons are not presynaptic to PAM neurons. These findings taken together suggest that all or some *DvPdf-*positive but *Pdf*-negative cells (i.e., the three CRY-negative LNds, one CRY/ITP double-positive LNd, and the fifth LNv) are presynaptic to PAM neurons (Fig. 4D).

### Identification of the vulnerable subpopulation of PAM neurons

PAM neurons are highly heterogeneous, encompassing 14 subclasses of morphologically and functionally diverse neurons projecting to different subdomains of the MB and other brain regions^38^. Because the number of neurons lost following H_2_O_2_ treatment was relatively consistent between experiments within the same genotype, we wondered if the vulnerable neurons belong to a specific subset(s) of PAM neurons. To test this possibility, we first used classical GAL4 drivers to label most or some PAM neurons with GFP to roughly locate the subclass vulnerable to H_2_O_2_ administered at ZT20. Most (8 of 13) PAM neurons that degenerated after the H_2_O_2_ insult belonged to the group expressing *R58E02-GAL4* (Fig. 5A).

**Fig. 5.**
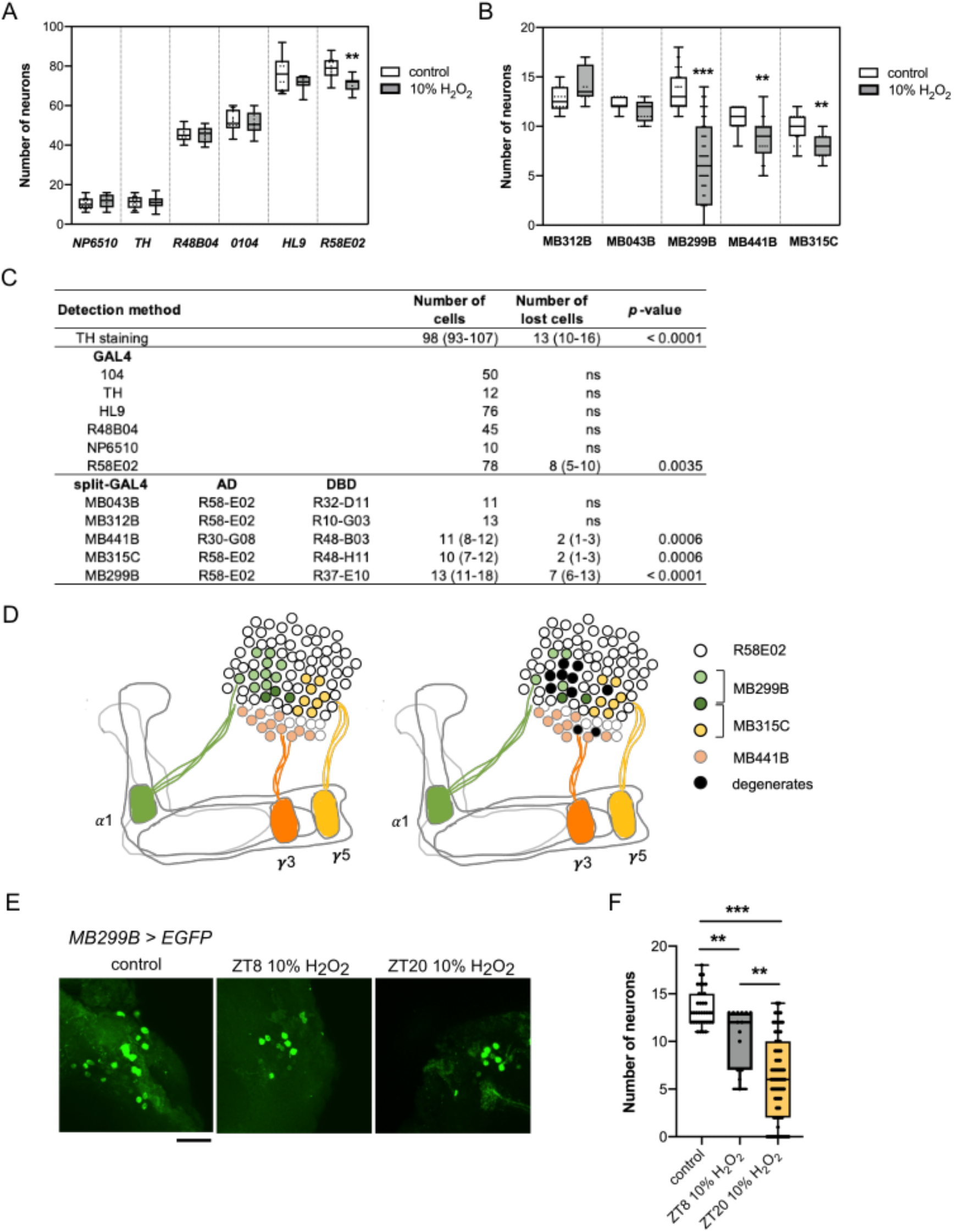
Identification of the vulnerable subpopulations of PAM neurons. (**A** and **B**) Quantification of the number of neurons labeled by GFP driven with different PAM neuron drivers, 7 days after a 4-h 10% H_2_O_2_ treatment at ZT20. The controls were treated with water only. n = 13–40 hemispheres. (**A**) *UAS-EGFP* was driven with classical *GAL4* drivers as indicated in the x-axis. Among the drivers tested, only the *R58E02*-labeled neurons include a subpopulation vulnerable to oxidative insults. \*\**p*<0.01 (t-test). (**B**) PAM subpopulations were labeled with split-*GAL4* drivers. Neurons labeled by MB299B, MB441B, and MB315C contain vulnerable PAM subpopulations. \*\**p*<0.01 and \*\*\**p*<0.001 (t-test). (**C**) Summary of the results from (**A**) and (**B**). The mean number of cells labeled by the given driver and of cells lost following the H_2_O_2_ treatment are shown. The numbers in the brackets indicate the range of values. AD, activation domain. DBD, DNA-binding domain. (**D**). Schematic representation of PAM subpopulations vulnerable to oxidative stress. PAM-*α*1, -*γ*5, and -*γ*3 neurons are labeled by MB299B, MB315C, and MB441B, respectively, with three PAM-*α*1 neurons co-expressing MB299B and MB315C (left). H_2_O_2_ treatment selectively degenerates approximately half of PAM-*α*1 neurons and a few cells each from PAM-*γ*5 and -*γ*3 subgroups (right). (**E** and **F**) 4-h 10% H_2_O_2_ treatment was performed at ZT8 and ZT20, and the number of MB299B-positive neurons was quantified 7 days post-treatment. (**E)** Representative confocal images of MB299B>EGFP neurons. Scale bar, 20 *μ*m. (**F**) Both experimental groups have significantly fewer remaining neurons than the control group treated with water only at ZT20. H_2_O_2_ treatment at ZT20 caused greater loss of neurons than at ZT8. \*\**p*<0.01 and \*\*\**p*<0.001 (t-test).

To identify the vulnerable PAM subgroup within the *R58E02*-positive cells, we tested split-GAL4s. From the available split-GAL4 drivers created at the Janelia Research Campus^39^, we selected all lines with activation domains driven with the *R58E02* promoter, i.e., MB312B, MB043B, MB299B, and MB315C. *UAS-GFP* was expressed with each of these drivers, and the number of remaining GFP-positive cells after short-term H_2_O_2_ treatment was scored. Neuronal loss was observed within the clusters labeled by the MB315C and MB299B drivers (Fig. 5B and C). PAM neuron subsets labeled by MB315C innervate the *γ*-lobe, and those expressing MB299B innervate the α-lobe of the MB^38^. Therefore, we tested one more split-GAL4 driver (MB441B) that does not contain the *R58E02*-driven component but is expressed in the PAM neuron subset projecting to the MB *γ*-lobe. As expected, H_2_O_2_ treatment caused a loss of neurons within the subpopulation labeled by MB441B (Fig. 5B and C). These results suggest that PAM neurons innervating the MB *γ*3, *γ*5, and α1 subdomains (PAM *γ*3, *γ*5, and α1 subgroups, respectively) are subpopulations vulnerable to neurodegeneration after oxidative insults (Fig. 5D). Of these, the PAM α1 subgroup labeled by MB299B was the most vulnerable, as more than 50% degenerated after the H_2_O_2_ treatment (Fig. 5B-D).

We next determined whether these PAM subpopulations were rhythmically vulnerable to oxidative stress, focusing on the most vulnerable MB299B-labeled neurons (PAM-α1). We treated 7-day-old flies expressing GFP with the MB299B split-GAL4 at ZT8 and ZT20 with 10% H_2_O_2_, and the GFP-positive cell count was scored 7 days later. The H_2_O_2_ treatment at ZT8 caused the loss of around three to four neurons, whereas, on average, 7 neurons were lost from the treatment at ZT20. Thus, PAM-α1 indeed showed the rhythmicity in vulnerability to oxidative stress (Fig. 5E and F). We also analyzed the Janelia Research Campus hemibrain connectome data^40^, which were visualized using the NeuPrint interface^41^ to assess the structural connectivity between clock neurons and MB299B neurons. Visual inspection of the s-LNvs, LNds, and MB299B neurons revealed close contacts between the axonal projections of the LNds and MB299B neurons (Fig. 6A), whereas projections of the s-LNvs and MB299B neurons did not show apparent contacts (Fig. S3A). These findings are consistent with the results of the *trans*-Tango experiment (Fig. 4C) and suggest that input from the LNds modulates the vulnerability of MB299B neurons in a circadian fashion. We found that the projections of the LNds also contacted MB315C and MB441 neurons, supporting the notion that vulnerable PAM neuron subsets are postsynaptic to the LNds (Fig. S3B and C).

**Fig. 6.**
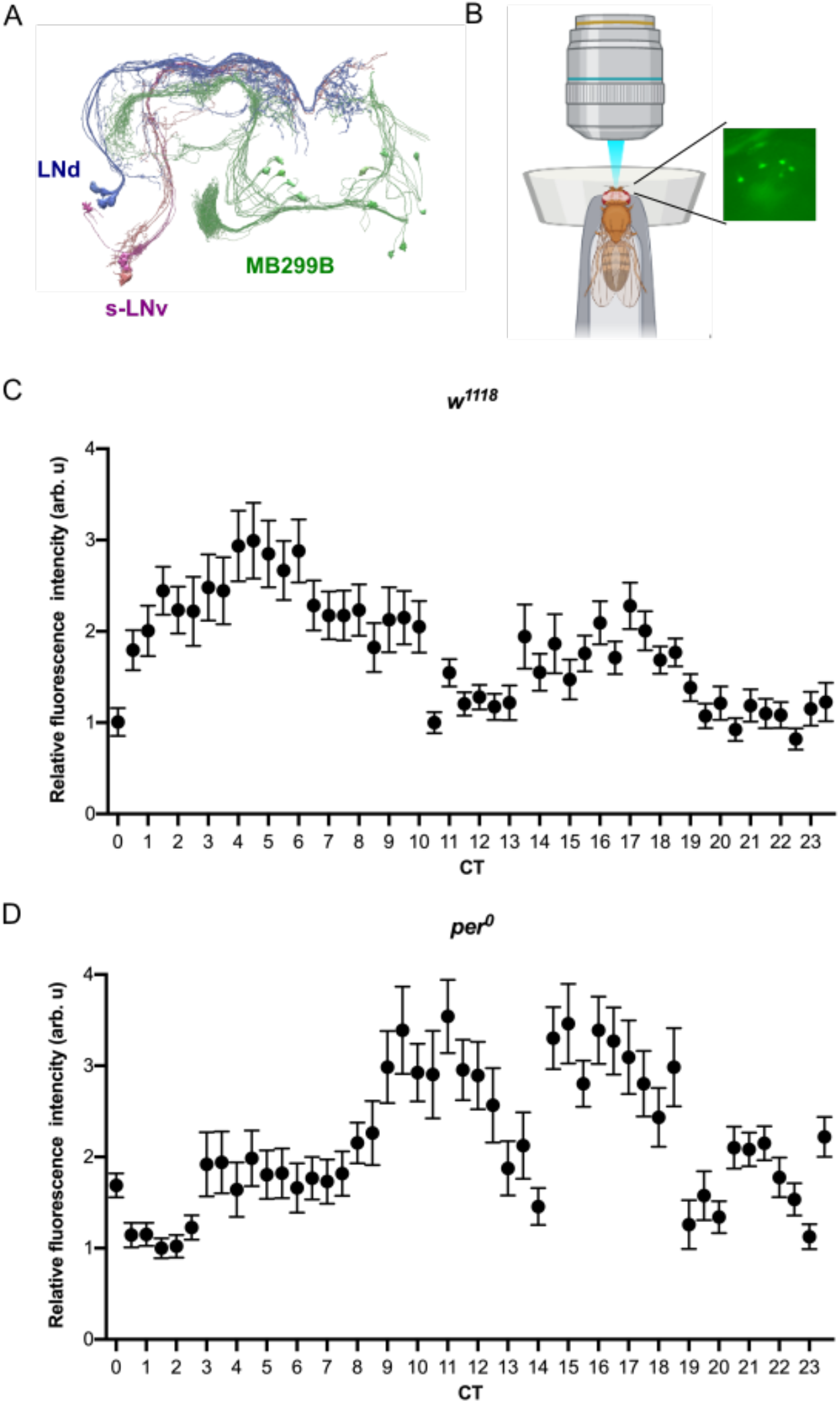
MB299B neurons display Ca^2+^rhythms. (**A**) Projections of the LNds contact with dendritic arbors of MB299B neurons. Image created using the hemibrain connectome data ^40^ via the NeuPrint tool ^41^. (**B**) Illustration of the method for live GCaMP imaging. (**C** and **D**) *MB299B>GCaMP7s* fluorescence levels in MB299B neurons throughout 24 h on the first day in DD following LD-entrainment in *w*^*1118*^ (**C**) and *per*^*0*^ (**D**). Relative fluorescence intensity (mean ± SEM) from 3 independent experiments. At each timepoint, n = 53–146 cells from 5–9 *w*^*1118*^ flies were analyzed in (**C**) and n = 22–112 cells from 2–8 *per*^*0*^ flies were analyzed in (**D**).

### Rhythmically vulnerable PAM subpopulations exhibit calcium rhythms

The finding that MB299B-positive, PAM-*α*1 neurons are rhythmically vulnerable to oxidative insults and postsynaptic to LNds raises the possibility that they exhibit physiological rhythms. Therefore, we measured levels of intracellular Ca^2+^, a second messenger that controls several cellular functions, in PAM-*α*1 neurons across 24 h in live flies (Fig. 6B). The calcium sensor GCaMP7s^42^ was expressed with the MB299B driver, and the 3-day-old flies were entrained to LD cycles for four days before imaging in DD. Ca^2+^ levels in these neurons showed temporal variations, with peaks twice a day, around CT5 and CT17 (Fig. 6C). In *per*^*0*^ flies, GCaMP fluorescence levels were uneven across timepoints and the bimodal pattern in *w*^*1118*^ was lost. These results suggest that circadian inputs to the PAM-*α*1 neurons rhythmically modulate their activity that peaks in the morning and early night.

Because Ca^2+^ levels in neuronal cell bodies can be correlated with firing frequency^43^, we wanted to know whether decreased excitability would be neuroprotective. Therefore, we silenced PAM neurons by overexpressing the inward rectifier K+ channel, Kir2.1^44^ with the *R58E02* driver and tested whether this silencing prevents degeneration of PAM neurons after the oxidative stress challenge. However, significantly fewer PAM neurons were present in the 14-day-old flies expressing Kir2.1 than in the control without H_2_O_2_ treatment, probably due to developmental impairments. Moreover, Kir2.1 overexpression did not prevent H_2_O_2_-induced PAM neuron loss (Fig. S4). This finding suggests that neuronal activity levels are unlikely to be the cause of the selective and rhythmic vulnerability of PAM-*α*1 neurons.

### Consequences of PAM-*α*1 neurodegeneration on non-motor functions

Four-hour treatment with 10% H_2_O_2_ at ZT20 leads to the loss of approximately thirteen DA neurons in the PAM cluster, most of which belong to the PAM-*α*1 subclass (Fig. 5A-D). We wondered whether this loss of approximately 10% of PAM neurons would have immediate or long-term effects on motor or non-motor functions. It was reported that synaptic dysfunction of a small subset of PAM neurons (expressing *NP6510-GAL4*) could impair the fly’s ability to climb^19^. To test whether the PAM-*α*1 subgroup has a similar role in climbing ability, we performed a startle-induced negative geotaxis assay^20^ after the short-term H_2_O_2_ treatment with increasing age. Climbing abilities of control and treated flies declined with age, as expected and as previously reported^19^. However, there was no difference between the control and experimental groups up to day 56, i.e., 49 days post-H_2_O_2_ treatment (Fig. S5), except for day 49. These findings suggest that PAM-α1 is not involved in the control of climbing behavior.

DA neurons play various roles in regulating behavior and physiology, including sleep. Dopaminergic input to the MB regulates several aspects of sleep^45^. Because the role of the MB *α*1 compartment on sleep remains elusive, we next examined the effect of H_2_O_2_-induced loss of PAM-*α*1 on sleep. Following the short-term H_2_O_2_ treatment on day 7 in LD, flies were maintained under LD cycles, and sleep was analyzed over three days starting at days 11 and 17. The results showed an increase in sleep in H_2_O_2_-treated flies in both age groups (Fig. 7A). Sleep increase occurred specifically during the night and was more pronounced in older flies (Fig. 7B and C). In contrast, activity levels of the flies during the waking period were not different between the H_2_O_2_-treated and non-treated groups at both ages (Fig. 7D and E). These findings suggest that PAM-*α*1 neurodegeneration caused by the H_2_O_2_ treatment increases sleep but not hypoactivity.

**Fig. 7.**
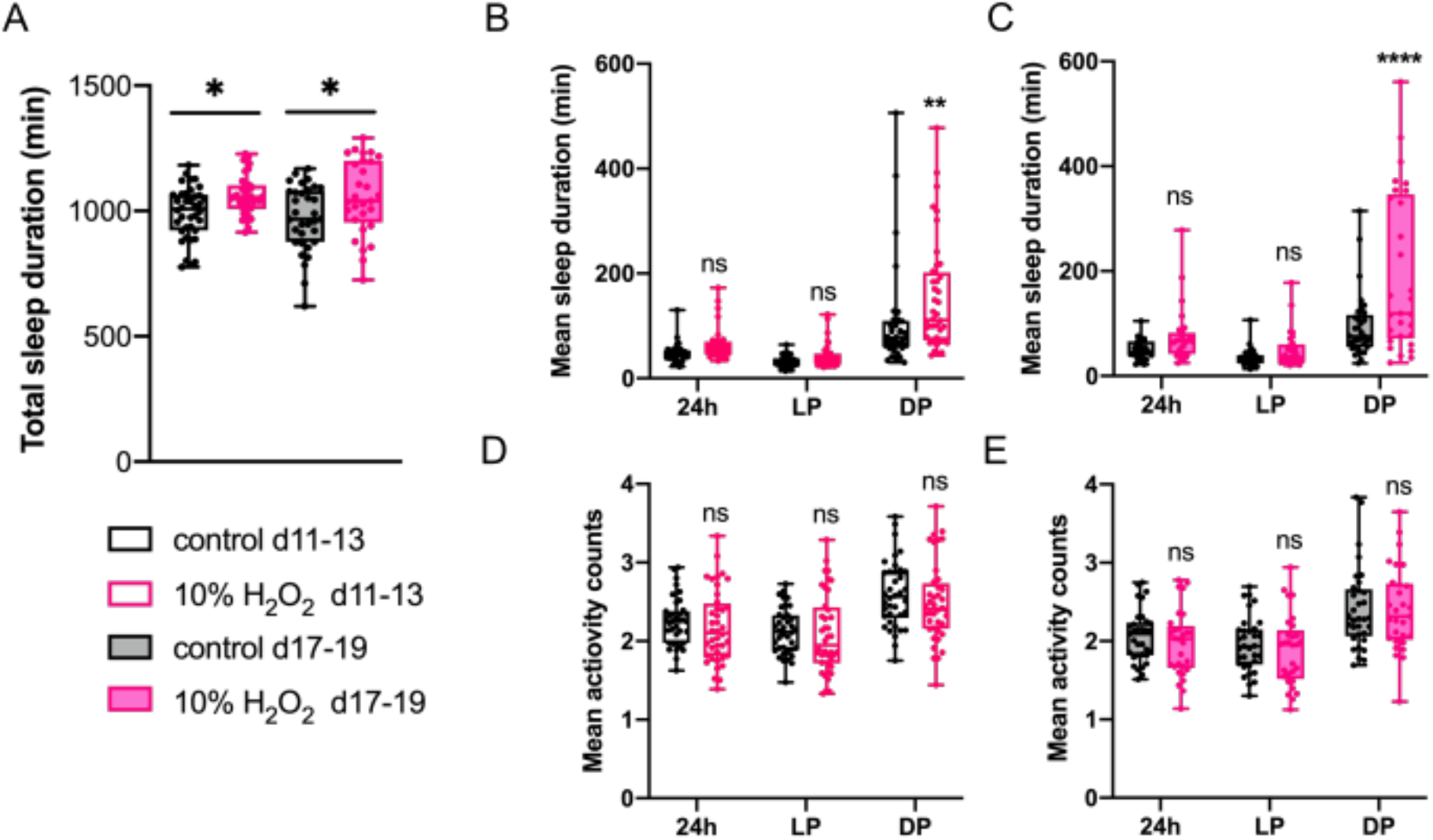
A single short-term H_2_O_2_ treatment increases nighttime sleep. 7-day-old *w*^*1118*^ flies were treated with 10% H_2_O_2_ or water for 4 h starting at ZT20 in LD and then placed in the activity monitor. The sleep and activity of flies from age day 11 to 13 (d11-13) and day 17 to 19 (d17-19) were analyzed. (**A**) Total sleep duration over 24 h in water-only control and H_2_O_2_-treated flies. H_2_O_2_-treated flies show an increase in total sleep in both age groups. \**p*<0.05 (ANOVA with Tukey’s multiple comparisons test) (**B** and **C**) Mean sleep episode duration over 24 h (24h), during daytime (light period, LP) and during the night (dark period, DP) from day 11 to 13 (**B**) and day 17 to 19 (**C**). H_2_O_2_-treated flies show an increase in nighttime sleep episode duration in both age groups. \*\**p<* 0.01 and \*\*\*\**p*<0.0001 (ANOVA with Šídák’s multiple comparisons test). (**D** and **E**) Mean activity counts during the wake period. No significant difference was observed between the control and treated groups.

### The implication in the multiple-hit hypothesis for Parkinson’s disease

PD arises from the interaction of genetic and environmental risk factors and age. This notion, formulated as the multiple-hit hypothesis^46^, suggests that short-term H_2_O_2_ treatment is merely a single “hit” to the dopaminergic system. To test this idea, we quantified the number of PAM neurons in the H_2_O_2_-treated flies over the course of aging. The difference between the control and treated groups was significant 7 days after H_2_O_2_ treatment and was maintained up to day 56 (49 days post-treatment), with no acceleration of neurodegeneration in either of the two groups (Fig. 8A).

**Fig. 8.**
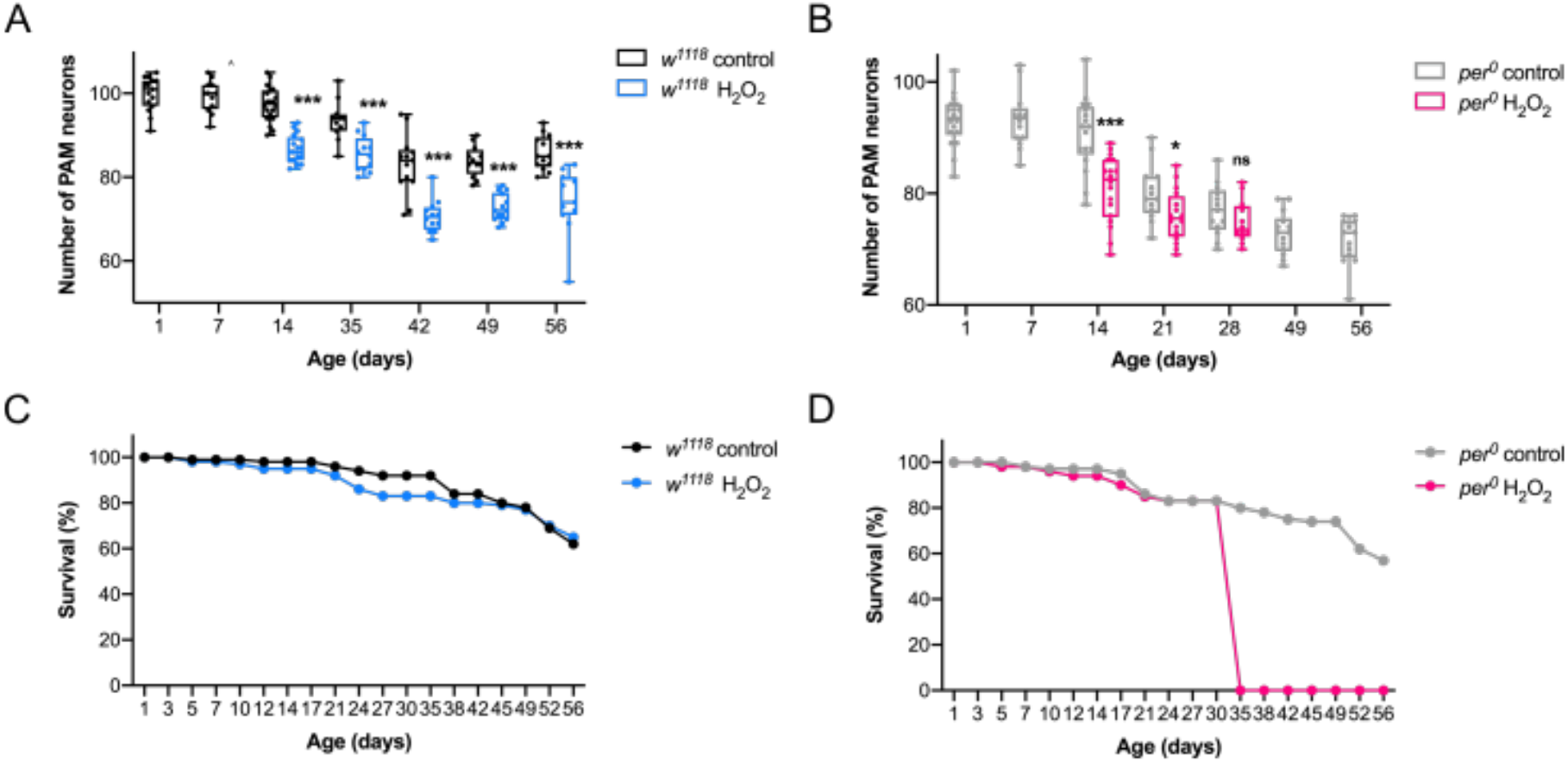
A single short-term H_2_O_2_ treatment causes premature death in *per*^*0*^flies. PAM neuron counts and survival of *w*^*1118*^ and *per*^*0*^ flies following a 4-h 10% H_2_O_2_ performed at ZT20 at 7 days old. (**A** and **B**) PAM neuron counts across aging in *w*^*1118*^ (**A**) and *per*^*0*^ (**B**). PAM neuron loss was observed as early as day 14 in both genotypes compared to control groups treated with water only. Neurodegeneration progresses with age but is not particularly accelerated in H_2_O_2_-treated flies. \**p*<0.05 and \*\*\**p*<0.001 (t-test). n = 12–20 hemispheres. (**C** and **D**) % of surviving flies in *w*^*1118*^ (**C**) and *per*^*0*^ (**D**). H_2_O_2_ treatment did not affect the survival of *w*^*1118*^ flies, but all H_2_O_2_-treated *per*^*0*^ flies died after day 30; in two independent experiments.

These findings support the multiple-hit hypothesis and suggest that an insult in addition to H_2_O_2_ might accelerate PAM neurodegeneration. We speculated that *per*^*0*^ mutation might constitute the second “hit,” following our finding that *per*^*0*^ mutants display age-dependent PAM neurodegeneration (Fig. 2A-C). Therefore, we counted the number of PAM neurons over the course of aging in *per*^*0*^ flies after the H_2_O_2_ treatment. On day 14, *per*^*0*^ showed a substantial loss of PAM neurons as expected; however, after that, the rate of PAM neurodegeneration in the H_2_O_2_-treated flies was not higher than in non-treated flies, reaching the same number of PAM neurons on day 28 (Fig. 8B). Strikingly, all the H_2_O_2_-treated *per0* flies died after day 28, whereas non-treated *per0*, as well as in H_2_O_2_-treated *w*^*1118*^ flies survived longer than 56 days (Fig. 8C and D).

Although after day 28, we could not determine whether the combination of H_2_O_2_ treatment and *per*^*0*^ mutation accelerated PAM neuron loss compared to *per*^*0*^ alone, these results nevertheless suggest that the combination of these factors causes premature death of the animal, reminiscent of increased mortality in PD patients^47^.

## Discussion

Circadian rhythm disruptions are frequent comorbidities in neurodegenerative disorders^48^; nevertheless, little is known about how circadian clocks and neurodegeneration are causally related. Here we showed that the circadian clock gene *per* controls daily rhythms and the magnitude of the vulnerability of DA neurons to oxidative insults in *Drosophila*. The circadian pacemaker circuit is presynaptic to the vulnerable subpopulation of DA neurons, PAM-*α*1, and rhythmically modulates its intracellular Ca^2+^ levels. Oxidative stress-induced dopaminergic neurodegeneration leads to persistent changes in sleep and, combined with the *per* null mutation, causes premature death of the animal. Taken together with the finding that *per* null mutation drives progressive loss of DA neurons, our results suggest a causal role of circadian clocks in dopaminergic neurodegeneration.

PAM-*α*1 neurons are highly vulnerable and rhythmically susceptible to oxidative stress. What makes this subpopulation more susceptible than other DA neurons and how circadian inputs gate the vulnerability remain questions for future studies. The LNd clock neurons are presynaptic to PAM-*α*1 neurons, and the latter exhibit *per*-dependent intracellular Ca^2+^ rhythms. Therefore, one can speculate that neuronal activity rhythms gated by the pacemaker circuit temporally regulate the vulnerability of PAM-*α*1. However, constant hyperpolarization by the expression of the Kir2.1 channel did not prevent H_2_O_2_-induced PAM neuron loss. The LNds express the neuropeptide ion transport peptide (ITP), the short neuropeptide F (sNPF), and acetylcholine (Ach)^49^. Whereas Ach is primarily excitatory, a synaptic structure that indicates fast neurotransmitter signaling was not detected between the LNds and PAM-*α*1 neurons in the connectome data^40^. Thus, the input from the LNds is likely neuromodulatory by the inhibitory sNPF^50^ or ITP, consistent with the inability of neuronal silencing to prevent PAM neuron loss. Identifying molecular differences between PAM-*α*1 and other subpopulations and potential molecular rhythmicity within PAM-*α*1 will be necessary to understand the mechanisms underlying these neurons’ selective and rhythmic vulnerability.

Ca^2+^ rhythms in MB299B-labeled neurons are bimodal, peaking around CT5 and CT17. This observation can be interpreted in two ways: all MB299B neurons exhibit bimodal Ca^2+^ rhythms, or there are two populations of neurons within the MB299B-labeled neurons that are anti-phasic to one another. Indeed, although PAM-*α*1 is the primary subtype labeled with the MB299B driver, weak expression of MB299B was also detected in PAM-*β*1 and *β*2 neurons^15,51^. Curiously, Ca^2+^ levels in the MB299B neurons in *per*^*0*^ show temporal variation, although its pattern is irregular and dissimilar to that of the *w*^*1118*^ flies. What causes this variation is unknown, but it might be an after-effect of the LD entrainment. Although further studies are required to untangle these issues, our observation is nevertheless consistent with the notion of connectivity between the circadian system and PAM neurons. Interestingly, the connectome data and results of the *trans*-Tango experiments suggest that the LNds are directly connected to the MB299B-labeled neurons but not the s-LNvs. However, restoring *per* expression within the PDF-positive LNvs is sufficient to reverse the PAM neuron loss in the basal conditions and under oxidative stress in *per*^*0*^ mutants. Because PDF-positive s-LNvs can synchronize the LNds through PDF/PDFR signaling^52^, these results suggest that the presence of PER within the PDF-positive s-LNvs is sufficient to convey a protective signal to the MB299B neurons via LNds.

Arrhythmic *per*^*0*^ and *tim*^*0*^ mutants display age-dependent loss of PAM neurons. Although both mutants completely disrupt molecular clocks, the observed phenotype could be a gene-specific effect rather than the result of loss of circadian clocks *per se*. A previous study demonstrated that *Clk* deficiency within the s-LNvs causes age-dependent loss of DA neurons in the PPL1 cluster and accelerates locomotor decline; however, the other clock gene loss-of-function mutants did not show the same phenotype^28^. Thus, the *Clk* has a role in preventing DA neuron loss and locomotor deficits in a circadian clock-independent manner^28^. Clock-independent roles of *tim* and *per* genes in mitochondrial uncoupling and lifespan have also been reported^53^. It is worth noting that MB299B neurons are not the first to degenerate in the basal conditions in *per*^*0*^ at least up to day 7, as these neurons are present at the age when Ca^2+^ imaging was performed. These findings underscore that clock genes play multiple roles in dopaminergic neuroprotection and are consistent with the notion that circadian disruptions are frequently comorbid with PD.

Short-term H_2_O_2_ treatment causes persistent increases in nighttime sleep but not locomotor defects, pointing to a role for PAM-*α*1 neurons in sleep regulation (but not excluding other PAM subpopulations). Lack of locomotor defects was expected, as another PAM neuron subset projecting to the MB *β*’ lobe is known to control startle-induced climbing ability^19^. It is well established that PAM-*α*1 neurons play an essential role in the acquisition and consolidation of appetitive long-term memory^15,54^ and the formation of courtship conditioning memory, which is a type of short-term memory^51^. Our findings that the circadian pacemaker circuit controls these neurons raise the possibility that these learning/memory performance types may show rhythmic variations across the day. Moreover, because they are highly susceptible to oxidative-stress-induced degeneration, their learning/memory functioning may be affected by oxidative stressors and are likely to be impaired when PAM-*α*1 undergoes degeneration. Thus, the circuity involving LNd – PAM-*α*1 – MB *α*1 subdomain has overlapping roles in sleep and learning/memory and is susceptible to oxidative stress. Impairments in this pathway might elicit phenotypes that are reminiscent of non-motor symptoms of PD.

Several lines of evidence support the multiple-hit hypothesis for PD, suggesting that progressive PD pathology is triggered by the interaction of multiple genetic/environmental risk factors^46^. Our finding that a single H_2_O_2_ treatment triggers DA neuron loss but that the rate of degeneration after that is not higher than the natural age-dependent loss in non-treated flies is consistent with the theory. We could not determine whether H_2_O_2_ treatment on *per*^*0*^ accelerates dopaminergic neurodegeneration, as this combination causes premature death. Of note, whereas immediate lethality has been reported when flies were treated with much lower concentrations of H_2_O_226_, we did not detect any such effect. Differences in a laboratory environment or fly strains might have caused this discrepancy. If not treated, PD presents a higher mortality risk, including sudden unexpected death^55^. Our findings are consistent with the multiple-hit hypothesis and suggest that genetic variations in circadian clock genes might represent a risk factor for PD. We also demonstrated that an oxidative stressor administered at a specific time of day could have a critical impact on the survival of DA neurons and cause persistent changes in behavior; these findings might also be relevant in humans.

## Methods

### Drosophila *strains and culture*

Flies were reared in 12-h LD cycles in a humidified chamber at 25 °C on a standard corn meal medium. The following lines were previously described: *HL9-GAL4* ^56^, *R58E02-GAL* ^14^, *per*^*0* 57^, *Pdf-GAL4* ^34^, *Clk1982-GAL4* ^33^, *DvPdf-GAL4* ^35^ *UAS-per16* ^58^. *NP6510-GAL4* (113956) was obtained from the Kyoto Stock Center. The following lines were obtained from the Bloomington Stock Center: *TH-GAL4* (Stock number 8848), *R48B04-GAL4* (50347), *0104-GAL4* (62639), *MB312B* (68314), *MB043B* (68304), *MB299B* (68310), *MB441B* (68251), *MB315C* (68316), *UAS-GCaMP7s* (79032), *UAS-Kir2*.*1* (6596), *UAS-myrGFP, QUAS-mtdTomato(3xHA)*, and *trans-Tango* (77124).

### H_2_O_2_ treatment

Male flies were collected after hatching and entrained to 12h/12h-LD cycles in an incubator at 25 °C. 7-day-old flies were transferred to an empty vial for 5 h, and then a filter paper soaked with 100 *μ*l H_2_O_2_ of a given concentration or H_2_O was inserted in the vial for a defined duration. Flies were then placed in a vial containing standard corn meal agar food for 7 days before the analysis of DA neuron integrity. For circadian H_2_O_2_ treatment in LD, newly hatched flies were entrained to LD cycles, and on day 7, flies were treated with 5% or 10% H_2_O_2_ for 4 h at one of the 6 ZT timepoints (ZT0, 4, 8, 12, 16 and 20) following a 5-h food and water deprivation. After the H_2_O_2_ exposure, flies were placed in the vial with the standard food and maintained under LD cycles at 25 °C for 7 days. For circadian treatment in DD, newly hatched flies were entrained to LD cycles for three days and then released to DD. After three DD cycles, flies were treated with H_2_O_2_ at one of the CT timepoints (CT0, 4, 8, 12, 16 and 20), as described above. Treatment and subsequent incubation were performed in DD.

### Immunohistochemistry

Immunostaining was performed on whole fly brains. Flies were decapitated, and the whole heads were fixed in 4% paraformaldehyde + 0.3% Triton X-100 on ice for 1 h. The heads were washed with 0.3% Triton X-100 in phosphate-buffered saline (PBT) three times at room temperature. The head cuticles were partly opened, and the heads were incubated in blocking solution (5% normal goat serum in PBT) for 1 h at room temperature. The incubation with primary antibodies was performed over two nights at 4 °C. Then, the heads were washed three times and incubated overnight with secondary antibodies at 4 °C. After three washes, the brains were dissected entirely by removing the remaining cuticle and trachea and ounted in Vectashield mounting medium (Vector laboratories, H-1000-10). The slides were stored at 4 °C and protected from light until imaging. The primary antibodies and the concentrations were as follows: rabbit polyclonal anti-TH (Millipore, ab152) 1:100; mouse monoclonal antibody nc82 (Developmental Studies Hybridoma Bank) 1:100; polyclonal rabbit anti-GFP (Invitrogen, A6455) 1:500; and monoclonal mouse anti-HA (Covance, MMS-101P) 1:200. The secondary antibodies and their concentrations were as follows: Alexa Fluor anti-rabbit 488 (Thermofisher, A21052) 1:250 and Alexa Fluor anti-mouse 633 (Thermofisher, A11008) 1:250.

### Feeding assay

The feeding assay was performed as previously described with minor modifications ^30^. Male flies were starved for 5 h before the assay (beginning of feeding). A filter paper soaked with a 0.125 mg/ml solution of Brilliant Blue FCF was then inserted in the vial for a control group. In the viral containing the flies of the experimental group, a filter paper soaked with a 0.125 mg/ml solution of Brilliant Blue FCF with 10% H_2_O_2_ was inserted. The filter paper was kept in the vials for 4 h, after which the flies were flash-frozen in liquid nitrogen. Then, the flies were vortexed to detach heads from bodies. A standard sieve (No. 25) was used to remove heads, and the bodies were transferred to 1.5-ml Eppendorf tubes. The bodies were homogenized with a motorized pestle in 1 ml of PBT and centrifuged at 13000 rpm for 15 min to clear the debris. The absorbance of the supernatant was measured at 625 nm on a spectrophotometer. The measured value was divided by the number of flies homogenized in one tube.

### ROS detection

MitoSOX Red (Invitrogen, M36008), a cell-permeable mitochondrial superoxide indicator, was used to measure endogenous ROS levels. Male flies were decapitated, and brains were dissected in Hanks’ Balanced Salt Solution (HBSS), followed by incubation in 5 *μ*M MitoSOX for 15 minutes at 37 °C and three washes with warm HBSS. The brains were mounted in HBSS and imaged immediately with Leica TCS SP5 confocal microscope.

### trans-Tango

The *trans*-Tango assay was performed as described previously^37^. Briefly, male flies were aged for 10 days at 18 °C to maximize *trans*-Tango expression with the optimal signal-to-noise ratio. The immunohistochemistry protocol described above was used to visualize the neurons and their projections.

### In vivo calcium imaging

The calcium sensor GCaMP7s^42^ was expressed with MB299B split-GAL4, and the 3-day-old flies were entrained to LD cycles for four days. On the first day in DD following the LD-entrainment, flies were placed under the fluorescence microscope in the perfusion chamber after the brief operation to remove a small part of the cuticle of the head capsule. Each experiment consisted of a session of 6-h to 8-h live imaging at 30-min intervals, starting at four different circadian times (CT4, CT10, CT16, and CT22). Images were acquired using the Leica DM550 fluorescent microscope. The exposed brains were constantly perfused with oxygenated saline to prevent desiccation or deterioration of the tissue during imaging. The fluorescence intensity of individual cells was measured using the freehand selection tool of Fiji/ImageJ. The corrected total cell fluorescence (CTCF) value was obtained by subtracting the measured fluorescence value outside of MB299B neurons from the measured fluorescence value inside MB299B neurons. The following formula was applied to calculate CTCF: CTCF = integrated density – (area of the region of interest x mean fluorescence of background). The mean of the CTCF of individual cells per timepoint was calculated, and the normalized CTCF time series was obtained by dividing the mean CTCF of each timepoint by the lowest mean CTCF over 24 h.

### Sleep assay and data analysis

The fly sleep assay was performed as described previously with minor modifications^24^. 7-day-old male flies were deprived of food and water for 5 h and treated with 10% H_2_O_2_ for 4 h starting at ZT20. The control group was treated with water only. After the treatment, flies were placed in individual glass tubes containing 5% agarose with 2% sucrose in the *Drosophila* Activity Monitoring System (Trikinetics) and entrained in 12 h:12 h-LD cycles at 25 °C for three days. Locomotor activity data were recorded during the subsequent 12 days in LD. We extracted sleep quantity from the activity data collected in 1-min bins (defined as bouts of inactivity lasting for 5 min or longer)^59^. Sleep analysis was performed in MATLAB (MathWorks, version R2017a) using SCAMP v2 (Sleep and Circadian Analysis MATLAB Program) according to the instruction manual (Vecsey laboratory)^60^.

### Startle-induced negative geotaxis assay

The startle-induced negative geotaxis assay was performed as previously described^61^ with minor modifications^20^. Twenty male flies were anesthetized with CO_2_ and placed in a 100-ml graduated glass cylinder. The cylinder was divided into five equal zones and graded from 1 to 5 from the bottom to the top. The bottom of the cylinder was marked as zone 0. After 1 h of recovery from the CO_2_ exposure, the climbing assay was performed at ZT2. Flies were gently tapped to the bottom of the column to startle them, from which they recovered quickly and started climbing the cylinder wall. The whole experiment was video recorded for subsequent analyses. The number of flies that climbed up to each zone within the 20 seconds after startling was manually calculated, from which climbing index (CI) was calculated using the following formula: CI = (0×n_0_ +1×n_1_ +2×n_2_ +3×n_3_ +4×n_4_ +5×n_5_) / n_total_, where n_total_ is the number of total flies used in the experiment and n_x_ is the number of flies that reached the zone X. Three trials for each experimental group were performed at 30-second intervals, from which the median values were chosen as final data of an experiment. Three to four independent experiments were performed per group per age.

### Confocal microscopy and image analysis

Fly brains were imaged using a Leica TCS SP5 confocal microscope. All images were analyzed using Fiji/ImageJ software (NIH). To count DA neurons, neurons positive for TH immunostaining or neurons labeled with fluorescent protein were counted manually using the cell-counter plugin of ImageJ through individual Z-stacks of the confocal images. To quantify the fluorescence intensity of the MitoSOX Red in the region of PAM neurons expressing GFP, the measurement feature of ImageJ was used after adjusting a proper threshold. To calculate the resulting fluorescence intensity of the MitoSOX Red in the region of PAM neurons, the CTCF value was obtained by subtracting the measured fluorescence value outside of the PAM region from the measured fluorescence value inside PAM neurons.

## Statistical analysis

GraphPad Prism (v.8.1) and R were used for statistical analysis and data plotting. All data were assessed using the D’Agostino-Pearson normality test. Normally distributed data were compared using parametric tests, and non-normally distributed data were analyzed using non-parametric tests. To compare two groups, the two-tailed Student’s t-test was used for normally distributed data, and for non-normally distributed data, the non-parametric Mann-Whitney test was used. To compare three or more groups of normally distributed data, one-way analysis of variance (ANOVA) with post hoc Tukey’s multiple comparison test and two-way ANOVA with Šídák’s multiple comparisons test were used. The Kruskal-Wallis one-way ANOVA with Dunn’s multiple comparison test was used to compare non-normally distributed data. The statistical significance threshold for all experiments was set at *p*<0.05. In all figures, stars represent statistical significance: * *p* <0.05, ** *p* <0.01, *** *p* <0.001 and \*\*\*\**p*<0.0001. ns indicates not significant.

## Acknowledgments

We thank Dong-gen Luo for helpful technical advice and the Bloomington Drosophila Stock Center and Kyoto Stock Center for fly strains. The following funding sources supported this research:

Swiss National Science Foundation grants (31003A_169548 and 310030_189169)

The Georges and Antoine Claraz Foundation

## Author contributions

Conceptualization: MMD and EN. Investigation: MMD, LCD, and EP. Visualization: MMD, LCD and EN. Writing: MMD, LCD and EN.

## Data and materials availability

All data are available in the main text or the supplementary materials.

