## Supplementary Materials for "Circadian clock disruption promotes the degeneration of dopaminergic neurons"

**This file includes:**

Figs. S1 to S5

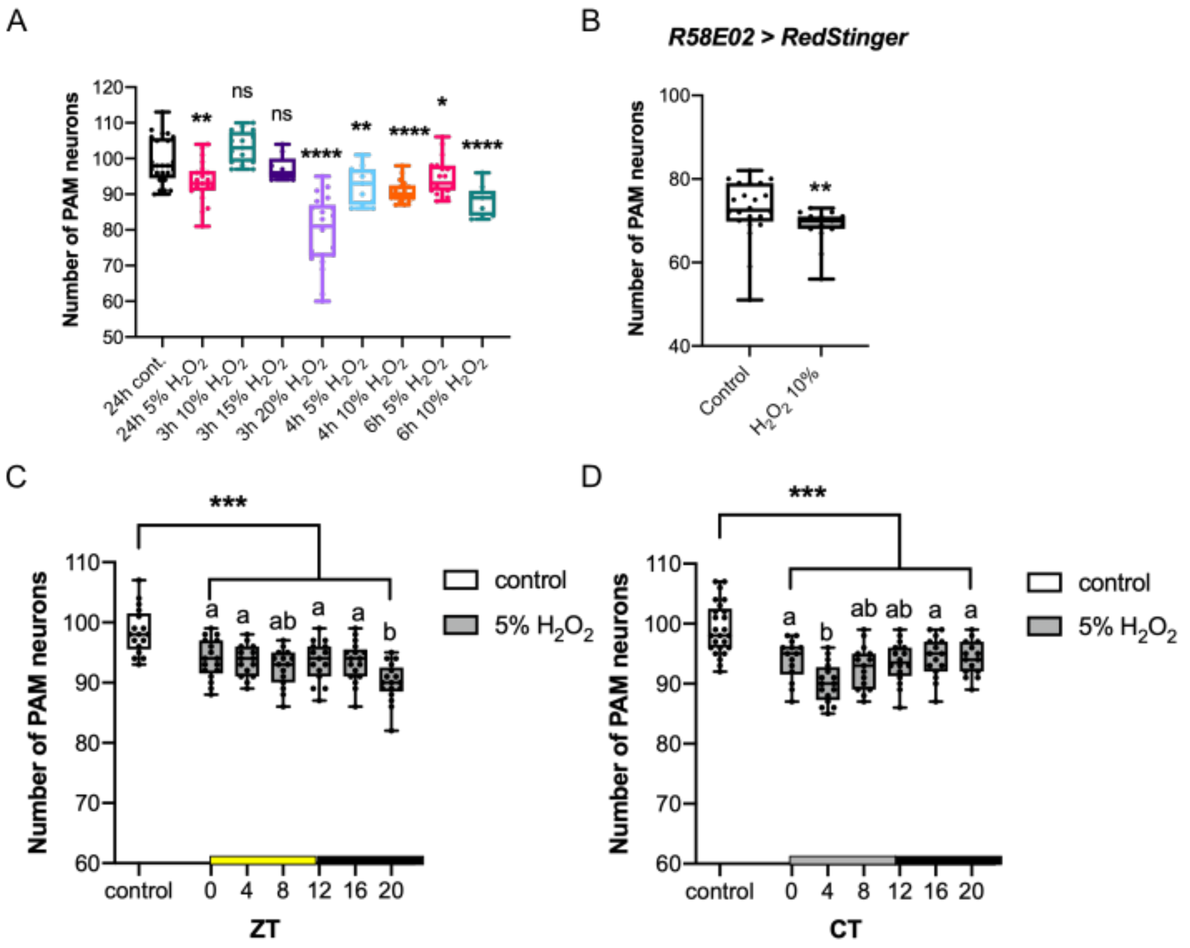


**Fig. S1. Short-term $H_2O_2$ treatment induces degeneration of PAM neurons**. (**A**) Effect of percentage and duration of $H_2O_2$ treatment on PAM neurodegeneration. 7-day-old $w^{1118}$ flies were treated with $H_2O_2$ of indicated doses, starting at ZT1 in LD. PAM neuron counts were examined by anti-TH immunostaining 7 days after the treatment. n = 10–24 hemispheres. *$p<0.05$, **$p<0.01$, and ****$p<0.0001$ by t-test comparing with the control treated with water only for 24 h (24 cont.). (**B**) PAM neurons were visualized with *UAS-RedStinger* driven by *R58E02-GAL4* and counted 7 days after a 4-h 10% $H_2O_2$ treatment of control treatment with water only. $H_2O_2$ treatment induces neuronal loss not just a reduction in TH levels. **$p<0.01$ (Mann-Whitney U test). n= 15–23 hemispheres. (**C** and **D**) PAM neuron counts were analyzed by anti-TH immunostaining 7 days after the 4-h 5% $H_2O_2$ treatment performed at different timepoints in LD (**C**) or in DD (**D**). The x-axis indicates the timepoints when $H_2O_2$ was applied. n=14-25 hemispheres. The control group was treated with water only at ZT20 in LD (**C**) and CT20 in DD (**D**). At all timepoints, PAM neuron counts in the $H_2O_2$ treatment group are significantly smaller than those in the control group. ***$p<0.001$ (ANOVA with Tukey's post-hoc test). Within the $H_2O_2$-treated group, flies treated at ZT20 in LD (**C**) and CT4 in DD (**D**) showed a significantly greater cell loss than the treatment at any other timepoint. Different lowercase letters represent statistical significance by ANOVA with Tukey's post-hoc test.

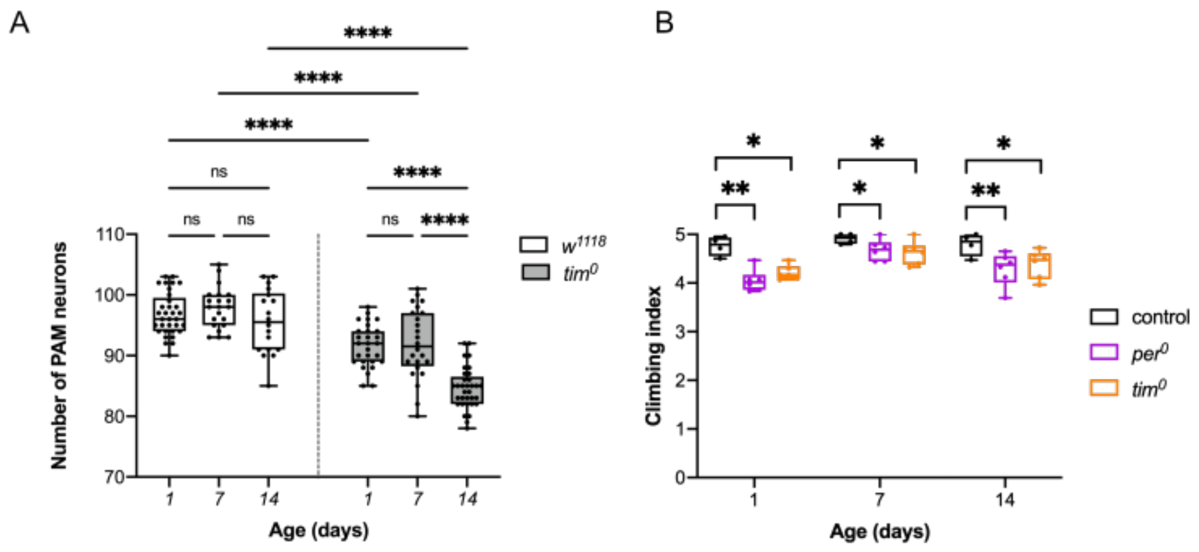


**Fig. S2. Progressive loss of PAM neurons in clock gene mutant flies.** (**A**) The number of PAM neurons in *w[1118]* and *tim[0]* at the indicated ages was analyzed by anti-TH immunostaining. *tim*[0] flies display developmental and age-dependent loss of PAM neurons. n = 18–33 hemispheres. ****$p$<0.0001 (ANOVA with Tukey's post-hoc test). (**B**) Negative geotaxis assays performed on *CS*, *per[0]*, and *tim[0]* flies at the indicated ages show climbing defects in clock gene mutant flies. 4–6 independent experiments. *$p$<0.05 and **$p$<0.01 (t-test with Welch correction).

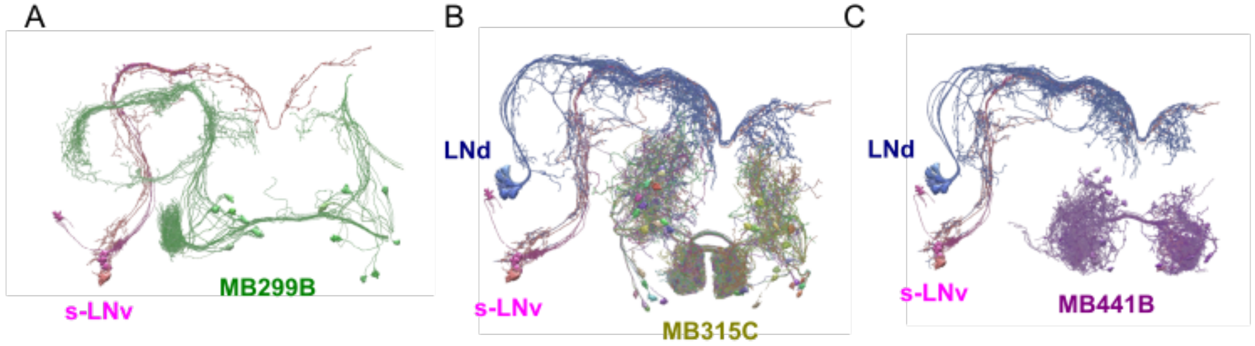


**Fig. S3. Position of PAM-$\alpha$1, -$\gamma$5 and -$\gamma$3 neurons and the s-LNv and LNd clock neurons.** (**A**) PAM-$\alpha$1 neurons labeled by the MB299B split-GAL4 driver and the s-LNvs do not contact. (**B**) Projections of PAM-$\gamma$5 neurons expressing MB315C contact the arbors of the LNds. (**C**) Projections of PAM-$\gamma$3 neurons labeled by MB441B do not contact the s-LNvs or the LNds. Images were created using the hemibrain connectome data with the NeuPrint tool.

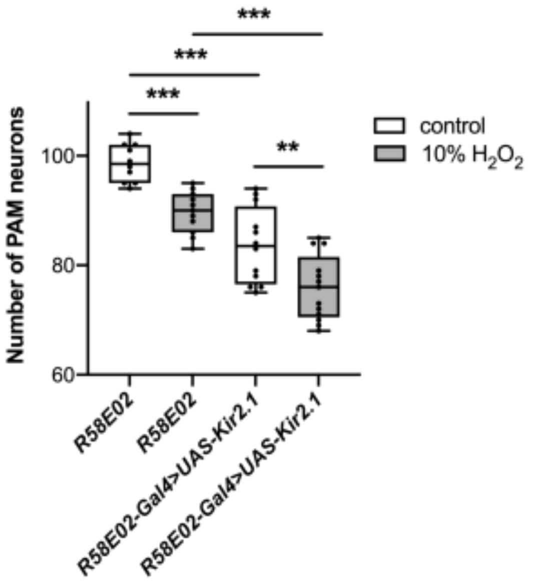


**Fig. S4. Effects of electrical silencing on PAM neuron development and degeneration.** Hyperpolarizing Kir2.1 channel was expressed in PAM neurons with the *R58E02* driver. PAM neurons were analyzed by anti-TH immunohistochemistry 7 days after a 4-h 10% $H_2O_2$ treatment performed at ZT20 or the control treatment with water. Kir2.1 expression did not prevent $H_2O_2$-induced PAM neuron loss and reduced the PAM neuron counts in basal conditions. n = 10–13 hemispheres. $^{**}p<0.01$ and $^{***}p<0.001$ (ANOVA with Tukey's post hoc test).

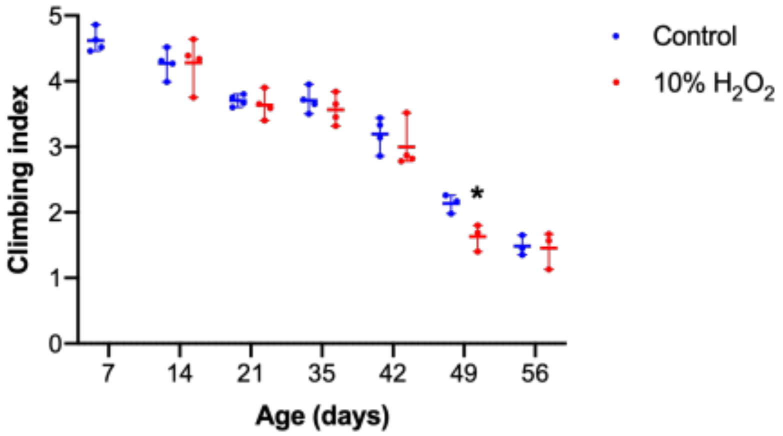


**Fig. S5. Short-term $H_2O_2$ treatment does not consistently impair climbing behavior.** The climbing behavior of the $w^{1118}$ flies was analyzed using the negative geotaxis assay following a 4-h 10% $H_2O_2$ or a control treatment performed at ZT20 at 7 days old; 4 independent experiments. Aside from day 49, no significant differences were observed between the control and the treatment groups throughout aging. *$p<0.05$ (t-test).
